# Microglia regulate nucleus accumbens synaptic development and circuit function underlying threat avoidance behaviors

**DOI:** 10.1101/2025.01.15.633068

**Authors:** Michael W. Gongwer, Fanny Etienne, Eric N. Moca, Megan S. Chappell, Sara V. Blagburn-Blanco, Jack P. Riley, Alexander S. Enos, Melody Haratian, Alex Qi, Rocio Rojo, Scott A. Wilke, Clare Pridans, Laura A. DeNardo, Lindsay M. De Biase

## Abstract

While CNS microglia have well-established roles in synapse pruning during neurodevelopment, only a few studies have identified roles for microglia in synapse formation. These studies focused on the cortex and primary sensory circuits during restricted developmental time periods, leaving substantial gaps in our understanding of the early developmental functions of microglia. Here we investigated how the absence of microglia impacts synaptic development in the nucleus accumbens (NAc), a region critical for emotional regulation and motivated behaviors and where dysfunction is implicated in psychiatric disorders that arise early in life. Using a genetically modified mouse that lacks microglia (*Csf1r* ^ΔFIRE/ΔFIRE^), we found blunted excitatory synapse formation in the NAc. This effect was most prominent during the second and third postnatal weeks, when we previously found microglia to be overproduced, and was accompanied by an increase in presynaptic release probability and alterations in postsynaptic kinetics. Tissue-level NAc proteomics confirmed that microglial absence impacted numerous proteins involved in synapse structure, trans-synaptic signaling, and pre-synaptic function. However, microglial absence did not perturb levels of astrocyte-derived cues and adhesive proteins that promote synaptogenesis, suggesting that reduced synapse number may be caused by absence of a microglial-derived synaptogenic cue. Although observed electrophysiological synaptic changes were largely normalized by adulthood, we identified lasting effects of microglial absence on threat avoidance behavior, and these behavioral effects were directly associated with alterations of NAc neuronal activity. Together, these results indicate a critical role for microglia in regulating the synaptic landscape of the developing NAc and in establishing functional circuits underlying adult behavioral repertoires.

## Introduction

The development of neural circuits relies on coordinated molecular and cellular interactions between neurons and glial cells. Key examples of these interactions include microglial regulation of axon guidance, regulation of programmed cell death, and regulation of synapse number via synaptic pruning^1–6^. A few studies have shown that microglia can also promote synapse formation and synapse maturation^7–9^, but these microglial contributions to synaptic development are much less studied. In addition, much of what we know about microglia in the developing brain comes from studies of the cortex and primary sensory circuits during restricted developmental time periods. This leaves substantial gaps in our understanding of microglial contributions to neural development.

The limbic system, including the nucleus accumbens (NAc), is important for emotional regulation and motivated behaviors^10^. The NAc mediates both reward approach and threat avoidance behaviors, and NAc dysfunction is implicated in a variety of psychiatric disorders that arise early in life^11,12^. However, our understanding of developmental trajectories of the NAc, and how they relate to subsequent behaviors, is lacking. Microglia have emerged as key regulators of synaptic development and are highly sensitive to environmental challenges that increase risk for psychiatric disorders. Given this, it is critical to understand how microglia shape synaptic development within limbic centers.

Maturation of neuronal populations and synaptic connections throughout the brain occurs in a highly coordinated, *region-specific* fashion^13,14^. We recently showed that microglial development is region-specific, with microglial density and morphology following distinct developmental trajectories in different regions of mesolimbic dopamine circuitry^15^. In the NAc, microglia displayed a robust overproduction during the second and third postnatal weeks, peaking at nearly double the density observed in the adult brain. In contrast, the ventral tegmental area (VTA) only exhibited a mild increase in microglial density during this period, suggesting a unique role for microglia in the developing NAc. The time course of NAc microglial overproduction did not align with programmed cell death or established periods of synaptic pruning, which occur earlier or later in development, respectively^15,16^. Studies in other striatal subregions indicate that the second and third postnatal weeks represent a period of pronounced synaptogenesis and refinement of synaptic kinetics^17,18^, suggesting that microglial overproduction during this time window may be related to their support of these processes within the NAc.

In this study, we used patch clamp electrophysiology, proteomics, and fiber photometry during behavioral assays in mice that possess or lack microglia to define potential roles for microglia in NAc synaptic development. We mapped the time course of synapse formation within the NAc and showed that the lack of microglia perturbs synapse formation during key developmental windows. To investigate potential molecules that drove those changes, we leveraged tissue proteomics and identified altered expression of microglial degradative enzymes and membrane proteins that may contribute to synapse formation. Although synaptic changes were less pronounced in adults, mice lacking microglia had altered activity in the NAc during a threat avoidance behavioral assay. Together these studies reveal how microglia regulate NAc developmental trajectories and adult behavioral repertoires and add to our understanding of region-specific roles for microglia during brain development.

## Results

### Absence of microglia blunts NAc excitatory synaptogenesis

We used a recently developed mouse line completely lacking microglia to investigate the impact of these cells on NAc synapse development. These mice were generated via deletion of fms-intronic regulatory enhancer (FIRE), a tissue-specific enhancer for colony stimulating factor 1 receptor (CSF1R), which microglia require for survival^19^. Immunostaining for microglia (Iba1/CD68) confirmed that prominent increases in NAc microglial density in the second and third postnatal weeks followed by a decrease by 3 months of age were observed in control (*Csf1r* ^+/+^, WT) mice from this line (Figure 1A-B), consistent with our previous findings^15^. In contrast, in NAc of *Csf1r* ^ΔFIRE/ΔFIRE^ (KO) mice, no Iba1+ or CD68+ cells were observed throughout early postnatal development or adulthood (Figure 1A-B).

**Figure 1:**
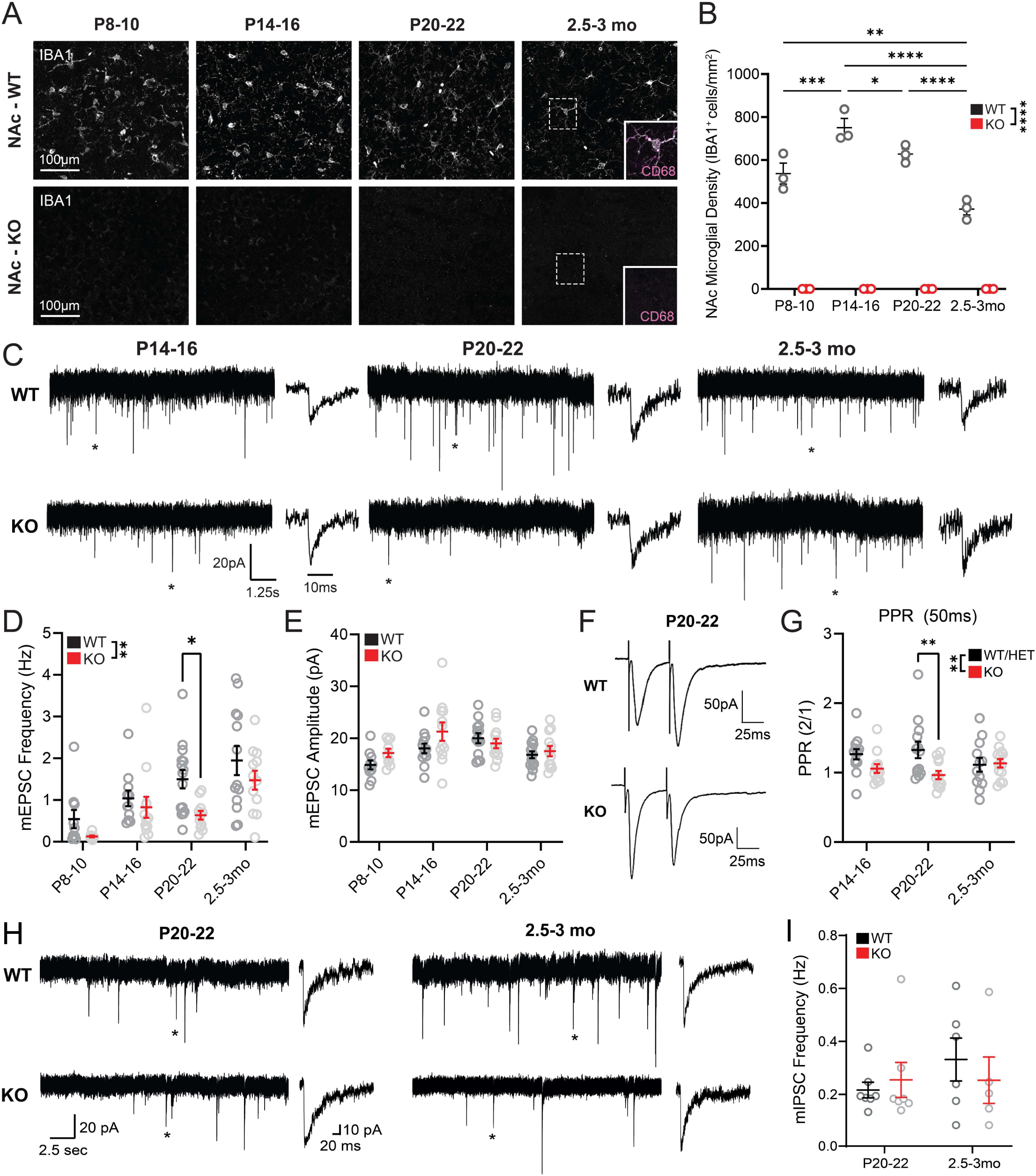
Absence of microglia blunts NAc excitatory synaptogenesis. (A) Representative images of NAc microglia in wild type (WT) and knockout (KO) mice at P8-10, P14-16, P20-22 and 2.5-3 months old. Microglia (IBA1) are represented in gray. Zoom panels represent the area with a dashed outline and also display CD68, a marker of microglial lysosomes, in magenta. Scale bar: 100 µm. (B) NAc microglial density across postnatal development in WT and KO mice. F_age_(3,16)=18.47 p<0.0001, F_genotype_(1,16)=947.0 p<0.0001, F_age x genotype_(3,16)=18.47 p<0.0001. Two-way RM ANOVA with Šídák’s multiple comparisons test. n=3 for each age and genotype. (C) Representative mEPSC traces in both WT and KO mice at p14-16, p20-22 and 2.5-3mo. Asterisks represent the single event shown for each trace. Cells were recorded while voltage clamping at -70mV in the presence of 100µM picrotoxin and 1µM TTX. (D) mEPSC frequency in NAc of WT and KO mice across postnatal development. F_age_(3,85)=11.01 p<0.0001, F_genotype_(1,85)=8.693 p=0.0041, F_age x genotype_(3,85)=0.7064 p=0.5508. (E) mEPSC amplitude in NAc of WT and KO mice across postnatal development. F_age_(3,86)=5.197 p=0.0017, F_genotype_(1,86)=3.064 p=0.0836, F_age x genotype_(3,85)=1.609 p=0.1932. P8-10: WT *n*=10(4), KO *n*=9(3); P14-16: WT *n*=12(6), KO *n*=12(5); P20-22: WT *n*=13(7), KO *n*=12(6); 2.5-3mo: WT *n*=14(7), KO *n*=12(6). (F) Representative traces of evoked paired-pulse ratio (PPR) recordings with a 50ms interstimulus interval in NAc knockout and control mice at P20-22. Cells were recorded while voltage clamping at -70mV in the presence of 100µM picrotoxin. (G) PPR for 50ms interstimulus intervals in NAc of KO and control mice at P14-16, P20-22, and 2.5-3mo. F_Age_(2,69)=0.1065, *P* =0.8991; F_Genotype_(1,69)=7.215, *P* =0.0090; F_Age x Genotype_(2,69)=2.669, *P* =0.0765. Two-way ANOVA with Šídák’s multiple comparisons test. (H) Representative traces of mIPSC events in both WT and KO at P20-22 and 2.5-3mo. Asterisks represent a single event shown on the right of each trace. Cells were recorded using high-chloride internal solution while voltage clamping at -70mV in the presence of 5µM CNQX and 1µM TTX. (I) mIPSC frequency in NAc of WT and KO mice at P20-22 and 2.5-3mo. F_age_(1,21)=0.7340 *P* =0.4013; F_genotype_(1,21)=0.09014, *P* =0.7669; F_age x genotype_(1,21)=0.7660, *P* =0.3914; \**P* <0.05, \*\**P* <0.01, \*\*\**P* <0.001, \*\*\*\**P* <0.0001. Error bars represent mean ± SEM.

To probe whether absence of microglia impacts excitatory synapse development in the NAc, we performed whole-cell patch clamp recordings from NAc medium spiny neurons in WT and KO mice at postnatal day (p)8-10, p14-16, p20-22, and 2.5-3 months (mo) of age (Figure 1C). These time points span the infantile to early juvenile period and young adulthood and capture periods when we observe prominent overproduction of microglia^15^. In WT mice, we observed a steady increase in mEPSC frequency across developmental time points but no changes in mEPSC amplitude (Figure 1D-E). Mice lacking microglia exhibited significantly lower mEPSC frequency across these timepoints, with this effect being most pronounced at p20-22. By adulthood, mEPSC frequencies in KO mice had increased, and did not differ significantly from WT levels. We also analyzed mEPSC kinetics at ages where these events were sufficiently abundant to allow rigorous fitting of average traces (Figure S1A). This revealed a significantly lower rise time in KO mice at p20-22 that remained significant in adulthood (Figure S1B), suggesting that there may be differences in AMPA receptor subunit composition at individual excitatory synapses of KO mice. No differences in mEPSC decay were observed at any age (Figure S1C).

A change in mEPSC frequency is typically driven by a change in either synapse number or synaptic release probability. To investigate potential changes in synaptic release probability, we performed paired-pulse electrical stimulation experiments in NAc at p14-16, p20-22, and 2.5-3 mo (Figure 1F, S1D). No significant changes in paired-pulse ratio (PPR) were observed across ages in WT mice (Figure 1G). However, PPR was significantly lower in KO mice compared to WT and heterozygous (*Csf1r* ^ΔFIRE/+^ HET) controls at p20-22 (Figures 1G, S1E), suggesting KOs have enhanced synaptic release probability. Therefore, reduced mEPSC frequency at this age is more likely to be driven by a reduction in synapse number. The elevated release probability in KO mice may represent a compensatory mechanism engaged in response to reduced excitatory synaptic input.

In the barrel cortex, microglia shape AMPA receptor insertion and abundance relative to NMDA receptors during postnatal development^20^. To further examine whether the absence of microglia affected the complement of AMPA and NMDA receptors at NAc synapses, we performed electrical stimulation and examined the AMPA / NMDA ratio by voltage clamping neurons at either -70mV or +40mV in the presence of picrotoxin, a GABA receptor antagonist (Figure S1F). The AMPA/NMDA ratios remained consistent across ages and genotypes (Figure S1G), suggesting that microglia do significantly impact relative abundance of these receptors at NAc glutamatergic synapses and further highlighting region-specific roles for microglia in supporting neural development.

### Absence of microglia alters NAc inhibitory synapse kinetics

Although microglial impact on synaptic function has largely been studied at glutamatergic excitatory synapses, microglia have also been shown to interact with inhibitory synapses in several brain regions^6,7,21^. To determine if the effects we observed were specific to excitatory synapses, we also recorded miniature inhibitory postsynaptic currents (mIPSCs) in p20-22 and adult mice (Figure 1H). No significant changes in mIPSC frequency or amplitude were observed across age or genotype (Figure 1I, S2A). However, significant differences were evident in mIPSC kinetics. mIPSC rise time was significantly faster in adult KO compared to WT mice, which was not observed at p20-22 (Figure S2B-C). KO mice displayed significantly higher mIPSC decay constant at p20-22 compared to WT mice, but this effect was no longer present in adulthood (Figure S2D). These findings raise the possibility that microglial absence is accompanied by differences in GABA_A_ receptor subunit composition and subtle differences in inhibitory synaptic input received by MSNs from KO mice in both development and adulthood.

### Absence of microglia alters developmental trajectories of astrocyte markers in NAc

Microglia and astrocytes can regulate each others’ properties^22–25^, and astrocytes can promote synapse formation and maturation^5,26–28^. To explore whether the absence of microglia affects NAc synaptic function indirectly via astrocytes, we immunostained for S100β, a calcium-binding protein expressed in astrocytes, and glial fibrillary acidic protein (GFAP), a marker of astrocyte reactivity (Figure S3A). In other brain regions, S100β and GFAP levels increase across postnatal development^29–31^. In line with this, the density of S100β+ cells in the NAc increased across the first three postnatal weeks in both WT/Het control mice and KO mice (Figure S3B). However, density of S100β+ cells was significantly higher in KO mice compared to WT/Het control mice at P20-22. In contrast to what has been observed in other brain regions, we observed a developmental decrease in the density of GFAP+ astrocytes in the NAc of WT/Het control mice (Figure S3C). This decrease was less prominent in KO mice, which had a significantly higher density of GFAP^+^ astrocytes compared to controls at both P20-22 and 2.5-3mo. These analyses indicate that the absence of microglia alters aspects of astrocyte development in the NAc, especially after the third postnatal week.

### NAc tissue proteomics highlights impact of microglial absence on synaptic function

Our electrophysiological measurements indicate that microglial absence impacts multiple aspects of NAc synaptic development and histological analyses raise the possibility that altered astrocyte development could contribute to synaptic changes. In particular, the prominent reduction in excitatory synapse number could be due to perturbation of known astrocyte synaptogenic molecules. To explore this possibility, we performed whole-tissue LC-MS/MS proteomic analysis on microdissected NAc from WT and KO mice at p20-22 (Figure 2A), when reduced mEPSC frequency is most prominent in KO mice. Differentially expressed proteins included, as expected, prominently downregulated microglia-specific (P2ry12, Hexb) and microglial-enriched (Ctsb, Ctsz, Npl, Cyfip1) proteins^32^ in tissue from KO mice (Figure 2B, Table S1). Moreover, proteins that are highly expressed in microglial-specific proteomic datasets (Myh9, Msn, Cap1)^33^ were also significantly downregulated in tissue from KO mice (Table S1), despite microglia only comprising roughly 5% of cells in the NAc. Proteins significantly upregulated in KO mice included GFAP, supporting histological observations that this astrocyte marker is elevated in the absence of microglia. However, astrocyte-derived synaptogenic cues that were detected (Sparcl1, Gpc4, Gpc6) did not differ significantly across genotypes (Figure 2C) and the majority of astrocyte-enriched proteins identified by astrocyte-specific proteomics were not significantly altered across genotype (Figure S4A). Families of adhesion molecules, such as neurexins, neuroligins, and protocadherins also play key roles in promoting synapse formation/maturation^34–36^. However, detected proteins within these families also did not differ significantly across genotype (Figure 2C). These observations suggest that the reduced number of functional excitatory synapses observed in KO mice via electrophysiology cannot be explained by insufficient abundance of astrocytic and adhesive proteins that promote synaptogenesis. Indeed, adhesion molecules Pcdhgc5 and Lrrtm3 were significantly *up*regulated in KO mice along with Snx32, a trafficking protein that regulates neurite outgrowth^37^, and Myo16, a myosin that regulates actin cytoskeleton reorganization^38^ (Figure 2B). Upregulation of these proteins may represent compensatory responses engaged by reduced synapse number.

**Figure 2:**
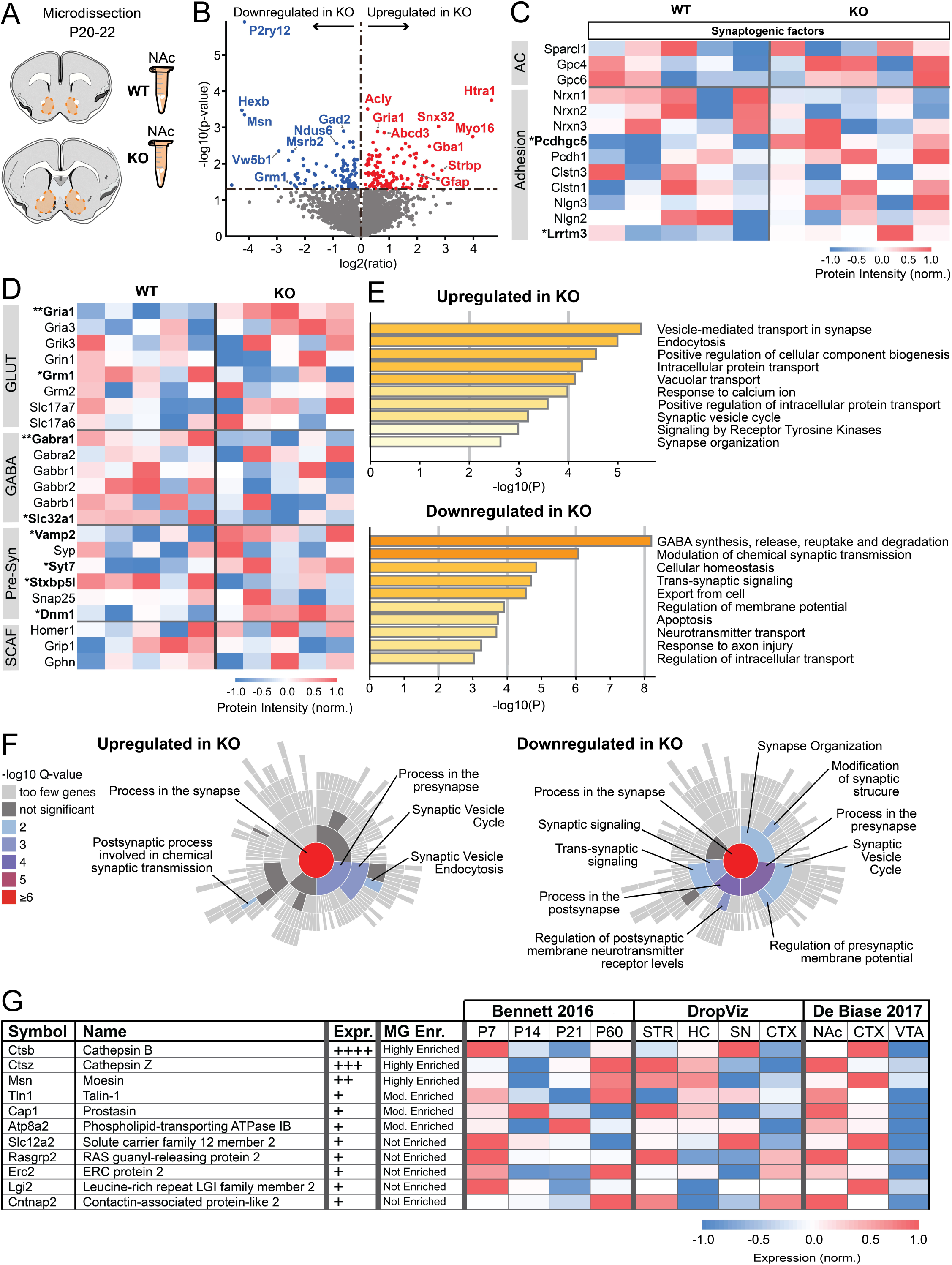
Microglia shape the developing NAc proteome. (A) Schematic representing NAc microdissection (orange) from acute brain slices at P20-22 in WT (*n*=5) and KO (*n*=5) mice. (B) Volcano plot representing up- and down-regulated proteins in NAc of KO mice compared to WT. (C) Heatmap representing relative expression levels of proteins known to be involved in synaptogenic cues. AC: astrocyte proteins. Asterisk represents proteins that are significantly up- or down-regulated in KO. *p<0.05. (D) Heatmap representing changes in protein expression involved in glutamatergic signaling (GLUT), GABAergic signaling (GABA), pre-synaptic signaling (Pre-Syn) and scaffold proteins (SCAFF). *p<0.05, **p<0.01. (E) Metascape pathway analysis of up- and downregulated proteins in KO mice compared to WT controls, showing top pathways relevant for molecular, cellular, or circuit-level information. Complete list of pathways is available in Figure S4. (F) SynGo synaptic protein enrichment analysis of up- and down-regulated proteins in KO mice compared to WT controls. (G) Table representing potential gene candidates extracted from three different published RNA sequencing datasets.

A reduced number of excitatory synapses could also be driven by deficits in essential protein components (i.e. the “building blocks”) necessary to assemble functional synapses. To determine if abundance of such proteins was reduced by microglial absence, we carried out additional, targeted analysis of key synaptic components. From a postsynaptic perspective, abundance of the majority of detected neurotransmitter receptors and scaffolding proteins did not differ across genotypes (Figure 2D). Consistent with our electrophysiological observations, upregulation of AMPA receptor subunit GluR1 (Gria1) in KO mice, could lead to more GluR2-lacking AMPA receptors with faster rise times^39^ (Figure S1B). Similarly, down-regulation of GABA_A_ receptor subunit ⍺1 (Gabra1), could lead to more ⍺2 containing receptors, which exhibit slower decay^40^ (Figure S2D). Pre-synaptic proteins involved in docking, fusion and recycling of synaptic vesicles, as well as neurotransmitter transporters, were either unaltered or showed a mix of up- (Vamp2, Syt7, Dnm1) and downregulation (Stxbp5l) across genotypes (Figure 2D). Upregulation of some of these proteins could contribute to alterations in release probability observed in our electrophysiology recordings (Figure 1F,G). Altogether, these observations suggest that, in general, protein components necessary to form functional synapses are still adequately produced in KO mice.

To explore proteomic results in a more unbiased manner and try to identify other potential mechanisms underlying reduced excitatory synapse number, we carried out pathway analysis of all significantly up- and downregulated proteins using Metascape^41^. Despite minimal changes in the *abundance* of core synaptic components and synaptogenic molecules, this analysis revealed alterations in multiple pathways associated with synaptic function, including synaptic vesicle recycling, neurotransmitter transport, and synapse organization (Figure 2E, S4B-C). Protein enrichment analysis with SynGo^42^ (Figure 2F), which probes for a large number of proteins that impact synapse function beyond simply neurotransmitter receptors, transporters, and scaffolding proteins, further supported the observation that microglial absence impacts numerous synapse-relevant proteins. Proteins involved in pre-synaptic function were upregulated in KO mice, while proteins involved in both pre-synaptic function as well as overall synapse structure and trans-synaptic signaling were downregulated in KO mice (Figure 2F). Collectively, these observations support the idea that changes in NAc synaptic function are a primary tissue-level effect of lacking microglia during postnatal development. Given that astrocytic and adhesive synaptogenic molecules were not significantly decreased by microglial absence, one possible explanation for reduced NAc synapse number in KO mice is that microglia directly promote NAc synapse formation or stabilization via cell surface or secreted molecules. To explore this possibility, we screened our proteomics dataset for proteins that were significantly downregulated in the absence of microglia and used functional annotation to further select proteins that are secreted or expressed on plasma membranes. Finally, we leveraged publicly available RNAseq datasets^32,43–45^ to define: 1) whether genes encoding those proteins are expressed at high or low levels in microglia, 2) whether expression is enriched in microglia compared to other CNS cells, 3) whether microglial expression is developmentally regulated, and 4) whether microglial expression is higher in striatum / NAc compared to other brain regions. This analysis pipeline identified multiple candidate molecules that microglia could use to promote excitatory synapse formation at this age (Figure 2G) and that warrant follow-up examination in future studies. Identified candidate molecules included secreted degradative enzymes Cathepsin B and Cathepsin Z, that could degrade extracellular matrix to permit synapse formation^46–48^, as well as Moesin, which regulates immunological synapse formation^49^, among other candidates.

### Absence of microglia perturbs avoidance behavior

Although electrophysiological measures of NAc synapse number were largely normalized by adulthood in mice lacking microglia, these changes could, nonetheless, produce lasting effects on behaviors that engage NAc circuits, such as threat avoidance behaviors^50–52^. To test this, we trained adult KO and WT mice in platform-mediated avoidance (PMA)^50,53^ (Figure 3A). In PMA, mice learn to associate a tone with a foot shock and that they can avoid impending shocks by escaping onto a safety platform. Both KO and control mice learned to avoid nearly all shocks by the end of the training session, but KO mice learned at a faster rate (Figure 3B). During a retrieval test one day later, tones were presented without shocks to examine memory-guided threat avoidance behavior (Figure 3C). When the tone played, control mice entered the safety platform with short latency and remained there during the final 2 seconds of the tone (when the shock would have occurred) on the majority of trials (Figure 3D-G). Surprisingly, KO mice displayed decreased levels of avoidance, marked by a longer latency to enter the safety platform, decreased time on the platform during tone periods, and fewer ‘successful’ trials (Figure 3D-G). We did not observe any differences in freezing behavior (Figure 3H), suggesting that differences in avoidance behavior did not arise from differences in the fearful association between the tone and the shock.

**Figure 3:**
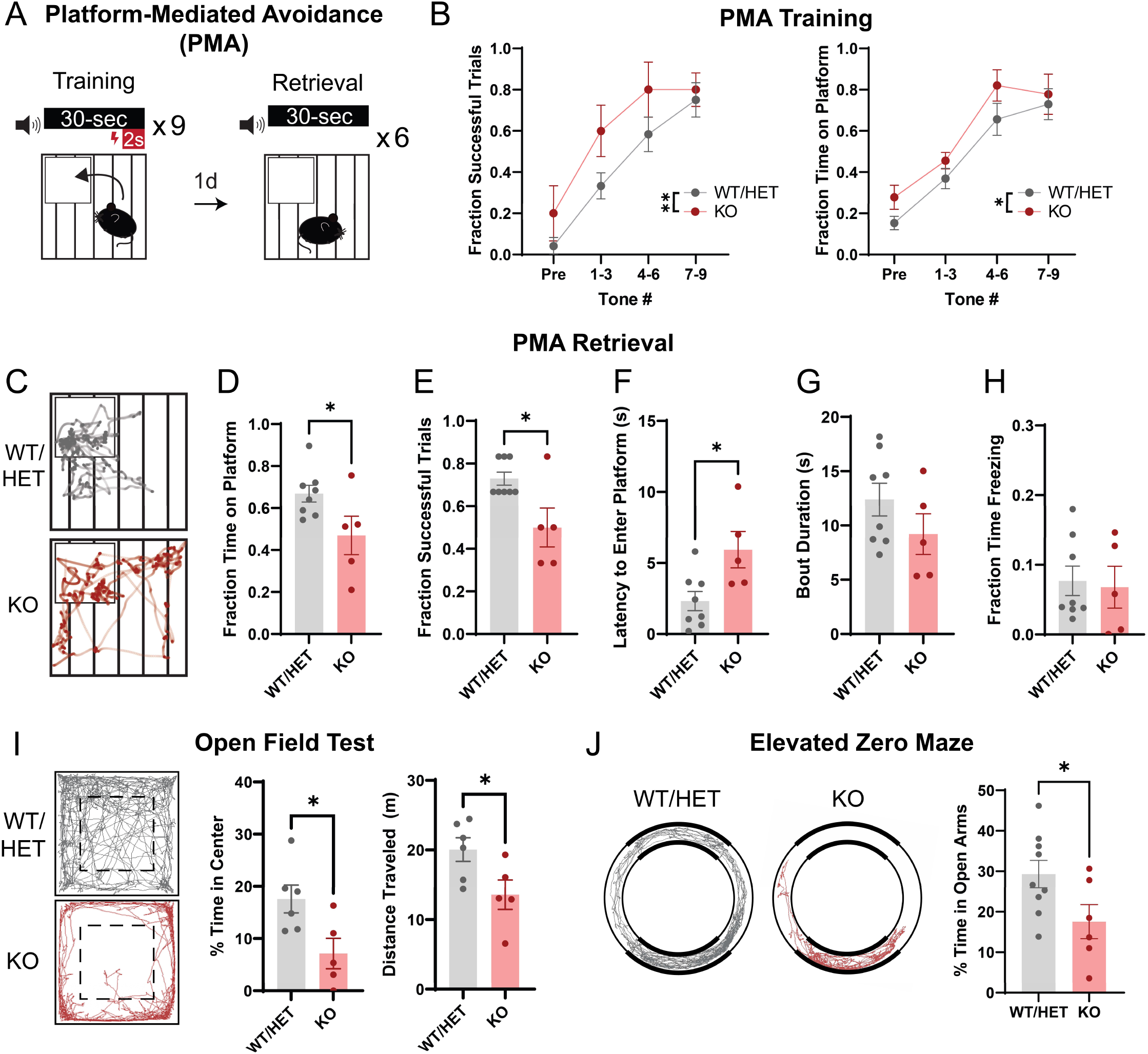
Absence of microglia disrupts learned and innate avoidance behavior in adulthood. (A) Schematic of behavioral paradigm. Mice received 9 tone-shock pairings on training day followed by 6 tones without shock on retrieval day. (B) PMA Training: Left, number of successful trials (i.e. successful avoidance of a foot shock) across training. F_Tone #_(3,33)=18.01, *P* <0.0001; F_Genotype_(1,11)=12.48, *P* =0.0047; F_Tone # x Genotype_(3,33)=0.4567, *P* =0.7143, Two-way RM ANOVA. Right, fraction of time spent on the safety platform during tone periods across training. F_Tone #_(3,33)=29.40, *P* <0.0001; F_Genotype_(1,11)=4.905, *P* =0.0488; F_Tone # x Genotype_(3,33)=0.2694, *P* =0.8470, Two-way RM ANOVA. WT/HET: *n*=8, KO: *n*=5. (C) Trajectory plots indicating the path of a representative mouse during tone periods of PMA retrieval. (D) Time on platform, (E) successful trials, (F) latency to enter the platform, (G) platform bout duration, and (H) freezing behavior during tone periods of PMA retrieval. Unpaired student’s t test, WT/HET: *n*=8, KO: *n*=5. (I) Behavior of KO and control mice in the open field test, including % time in the center of the arena and total distance traveled. Unpaired student’s t test, \**P* <0.05. WT/HET: *n*=6, KO: *n*=5. (J) Behavior of KO and control mice in the elevated zero maze. Unpaired student’s t test, \**P* <0.05. WT/HET: *n*=9, KO: *n*=6. \**P* <0.05, \*\**P* <0.01. Error bars represent mean ± SEM.

To determine if differences in PMA behavior were driven by differences in anxiety-like behaviors, we compared WT and KO mouse behaviors in the open field test (OFT) and the elevated zero maze (EZM). Mice innately avoid the center of an open field arena and the open arms of the EZM, as these are associated with higher levels of potential threat exposure. KO mice spent significantly less time in the center and travelled less distance in the OFT, and also spent significantly less time in the open arms of the EZM (Figure 3I-J). Hence, KO mice exhibit prominent anxiety-like phenotypes, suggesting that the decreased avoidance during PMA retrieval is driven by differences in learning and memory and not by lower levels of innate avoidance. We also compared sensitivity to the foot shock stimulus across WT and KO mice. KO mice generally had a higher shock threshold to elicit behavioral responses (Figure S5A), but a scurry response was observed in all mice well below 0.13 mA, the shock intensity used for PMA, suggesting that differences in shock sensitivity do not explain the behavioral differences we observed in PMA.

### Absence of microglia alters NAc activity during PMA

KO mice lack microglia throughout the CNS and microglial absence could impact neuronal function in multiple brain regions involved in successful PMA learning and memory. To probe whether changes in threat avoidance behavior specifically involve changes in NAc neuronal activity in KO mice, we used fiber photometry to record fluorescence activity from a virally-delivered, genetically encoded calcium sensor (AAV-syn-jGCaMP7f-WPRE) expressed in NAc neurons while mice performed PMA (Figure 4A). During training, foot shocks induced a brief, sharp increase in NAc activity in WT mice that rapidly decayed below the pre-shock baseline (Figure 4B). KO mice exhibited a similar peak compared to WT mice, but NAc activity remained elevated well after the shock, resulting in a significantly higher area under the curve (AUC) for the 20 second period post-shock (Figure 4B). During the retrieval test, we observed no notable fluctuations in NAc activity at the onset of the tone in both WT and KO mice, consistent with our previous reports^54^ (Figure 4C). During platform entries however, we observed an increase in NAc activity in WT mice that was absent in KO mice (Figure 4D). As an independent means of probing NAc activity levels in relation to behavior, we perfused mice 90 minutes after completion of PMA retrieval and immunostained for Fos, a marker of recently active neurons. KO mice had significantly fewer Fos+ NAc neurons compared to WT mice (Figure 4E), further demonstrating that the observed behavioral effects are associated with decreased NAc activity.

**Figure 4:**
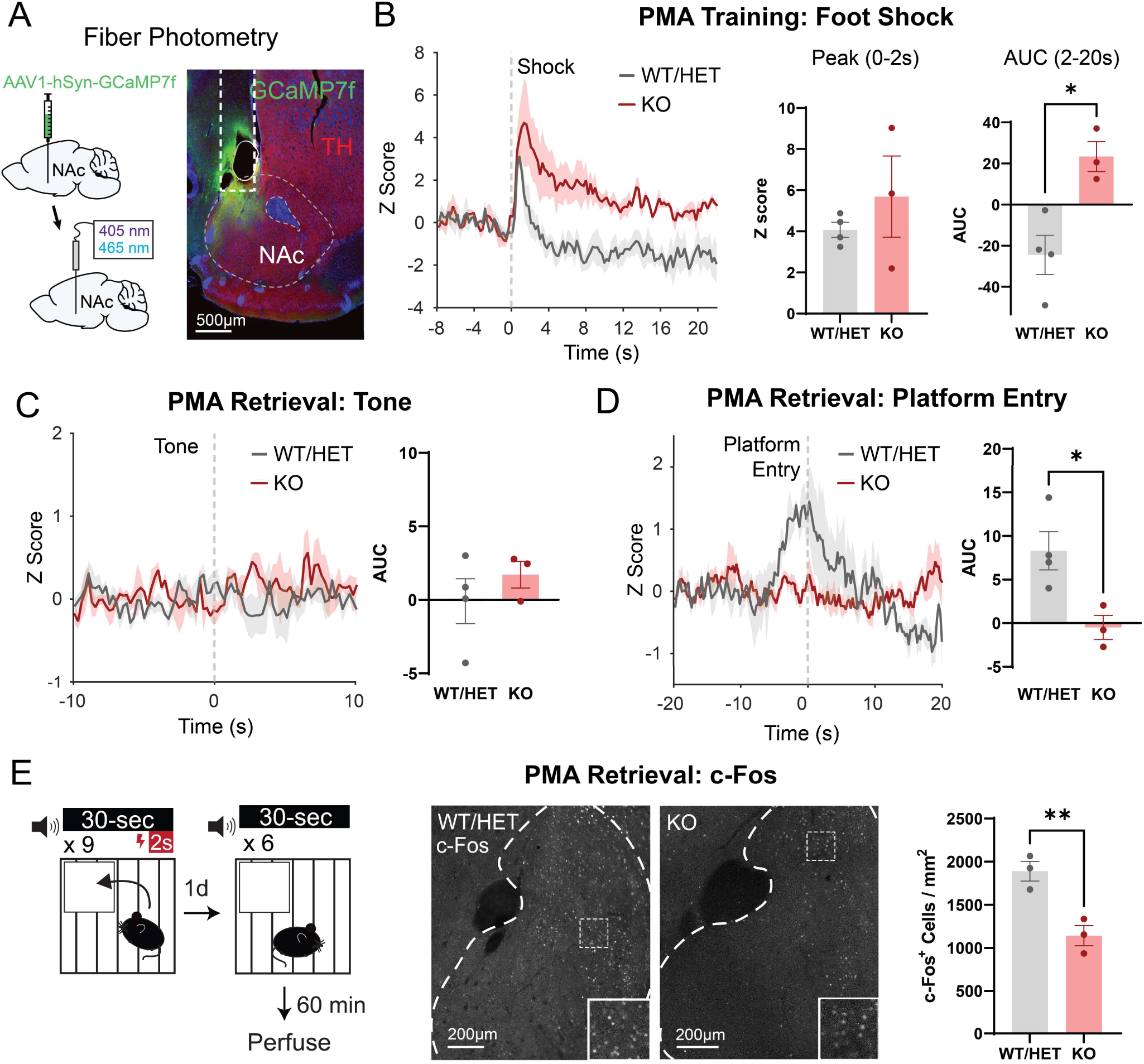
Absence of microglia alters *in vivo* function of the adult NAc during learned avoidance. (A) Injection strategy (left) and representative histological image (right) demonstrating viral expression and fiber placement in NAc. Scale bar 500µm. (B) Left, average traces representing foot-shock elicited NAc activity during PMA training in knockout and control mice. Error shading represents mean ± SEM by animal. Middle, peak shock response during the first two seconds following shock onset. Right, area under the curve (AUC) following the shock period. Unpaired student’s t test. WT/HET: *n*=4, KO: *n*=3 mice. (C) NAc activity at tone onset during the retrieval session (AUC: 0 to 10 seconds). Unpaired student’s t test. WT/HET: *n*=4, KO: *n*=3 mice. (D) NAc activity surrounding platform entry during the retrieval session (AUC: -10 to 5 seconds). Unpaired student’s t test. WT/HET: *n*=4, KO: *n*=3 mice. (E) c-Fos_+_ neuron density in NAc following PMA in knockout and control mice. Unpaired student’s t test. WT/HET: *n*=3, KO: *n*=3 mice. Scale bars 200µm. \**P* <0.05, \*\**P* <0.01. Error bars represent mean ± SEM.

## Discussion

Here we used the FIRE mouse line to study how the absence of microglia impacts the functional maturation of the NAc. Patch clamp studies provided key insights into the normative developmental trajectory of NAc synapse development, and revealed that FIRE mice had blunted excitatory synaptogenesis and increased synaptic release probability in juvenile stages of development (Figure 1). Proteomic analysis revealed that changes to synapse-relevant proteins is a primary tissue-level effect of lacking microglia in the developing NAc. Yet, astrocyte-derived synaptogenic cues and adhesion proteins that promote synapse maturation and stability were not affected, suggesting that lack of microglial-derived cues may underlie observed deficits in synapse formation (Figure 2). Using a combination of behavioral assays and fiber photometry, we found that in adult KO mice, disruptions in threat avoidance behavior were associated with changes in NAc activity (Figure 3,4). Together, our findings revealed that the absence of microglia lead to transient changes in NAc synapse development and lasting changes in NAc circuit activity and related behaviors. Moreover, we reveal key candidate proteins that may mediate synaptic changes in juvenile mice.

Although microglial removal of synapses via phagocytic engulfment has received the vast majority of attention when investigating synaptic interactions of these cells^55,56^, microglia also promote synapse formation and synapse maturation^7–9^. In the developing cortex (p8-10), microglia can induce dendritic spine formation by contacting neuronal dendrites and inducing local dendritic Ca^2+^ transients and actin remodeling^8^. Microglial depletion also reduces dendritic spines on immature, adult born granule cells in the olfactory bulb^57^, further supporting links between microglia and formation of post-synaptic structures at excitatory synapses. Microglia may play similar roles at inhibitory synapses; they promote formation of GABAergic axo-axonal synapses between chandelier cells and pyramidal neurons in cortex and these synapses are reduced by microglial depletion, microglial responses to LPS, or deletion of microglial GABA_B1_ receptors^7^. Perturbing microglial maturation via deletion of Cx3cr1 (fractalkine receptor) also indicates roles for microglia in synapse development, with Cx3CR1 KO mice displaying deficient AMPA receptor insertion and synapse maturation in early postnatal barrel cortex^20^ and reduced spine density in adult born granule cells in the olfactory bulb^57^. Despite multiple examples of microglial influence on synapse formation, microglial involvement in synaptogenesis has not been investigated in most brain regions and the molecular mechanisms underlying these microglial actions remain obscure.

Our results suggest that microglia influence multiple aspects of synaptic function in the developing NAc. We observed blunted NAc excitatory synaptogenesis throughout the first three postnatal weeks, with effects being most prominent in the third postnatal week (Figure 1). The first three postnatal weeks are a highly active period of synaptogenesis in dorsal striatum, and our results in WT mice similarly show increasing mEPSC frequency in the NAc throughout this period. Given our previous finding that NAc microglia are overproduced in the second and third postnatal week^15^, and the deficits in NAc synaptogenesis in KO mice, these findings raise the possibility that striatal microglia are specifically overproduced during this developmental window to support synapse formation. Our proteomics data pointed to the degradative enzymes Cathepsin B and Cathepsin Z as potential secreted microglial molecules that could regulate synapse formation. These cysteine proteases are highly expressed by microglia, including those in the dorsal striatum and NAc (Figure 2). Moreover, secreted cathepsins could support formation of additional synapses via degradation of extracellular matrix (ECM) and/or activation of growth factors such as BDNF^47,48^. Indeed, in other studies, we found that adult FIRE mice have elevated ECM accumulation in the NAc compared to control mice^58^, supporting the idea of microglial-based ECM regulation in this brain region. Microglial proteins that could play a role in contact-based promotion of synapse formation include Moesin and Talin-1. These molecules are enriched in striatal and NAc microglia and play key roles in cell-surface localization of integrins and formation of cell-cell focal adhesions^49,59^. Integrins are highly implicated in synapse formation^60^, raising the possibility that microglia stabilize nascent synapses via talin/moesin guided, integrin-based interactions with synaptic compartments. Thus, we identified key signaling proteins that may play previously unappreciated roles in microglia-mediated synapse formation in the NAc and that merit dedicated follow-up studies.

One limitation of whole-tissue proteomics is that these data cannot reveal whether some synapse-relevant proteins with normal abundance nonetheless have perturbed protein trafficking. Our proteomics pathway analysis indicated multiple hits related to intracellular protein trafficking (Figure 2), suggesting that approaches to investigate protein trafficking, particularly within neuronal dendrites and axons, should be a key future direction. Also, it is possible that decreased protein expression in one cell type could be masked by increased expression in another cell type. Future studies using cell-specific and/or cell compartment specific proteomics can be used to further explore the molecular mechanisms of how microglial absence impacts synapse development. Astrocytes also regulate synapse formation and function^5,26–28^ and we observed increases in multiple astrocyte proteins (Gfap, Htra1, S1pr1) in KO mice (Figure 2, S4), which may be driven by lack of microglia-derived regulatory cues or represent a compensatory mechanism. We did not observe changes in detected astrocyte-derived synaptogenic cues (Gpc4,

Gpc6, Sparcl1), suggesting that microglia are primary mediators of the synaptic changes we observed. However, some astrocyte-derived synaptogenic cues were not detected via proteomics (Thbs1, Thbs2, Sparc, Chrdl1) and we cannot exclude that these factors are present in the developing NAc at low levels and are perturbed by microglial absence.

Many of the effects we observed on functional synapse maturation in KO mice normalized to near-WT levels by adulthood, suggesting that compensatory synaptogenic mechanisms are engaged in other cell types between P22 and adulthood. Another possibility is that the juvenile-specific synaptic changes we observed may represent a developmental delay induced by chronic microglial absence, with ongoing synaptogenic mechanisms eventually able to reach near-WT adult levels. However, this eventual normalization of synapse number does not necessarily mean that specific patterns of circuit connectivity and circuit function are comparable across genotypes. Indeed, we observed lasting effects of microglial absence on NAc circuit function, with adult FIRE mice exhibiting anxiety-like behaviors, and impaired avoidance memory retrieval that was accompanied by lower levels of behavior-associated NAc activity (Figure 3-4). These findings suggest that shifts in the developmental wiring and/or developmental trajectory of the NAc may have permanently altered the ability of NAc neurons to contribute to threat avoidance behavior. An alternative possibility is that acute microglial absence in adulthood may interfere with normal mechanisms for synaptic plasticity and memory consolidation. In support of this idea, microglia in adult motor cortex contribute to motor learning by promoting synapse formation^9^. Finally, microglia also regulate neuronal excitability^61^ and our proteomics pathway analysis showed changes in membrane potential-regulating proteins (Figure 2). Thus, lasting changes in neuronal excitability could also contribute to the observed changes in NAc activity in adult FIRE mice (Figure 4). Additional studies are needed to investigate these possibilities.

Our findings contrast sharply with recent work showing normal synapse development in the hippocampus and barrel cortex of FIRE mice^62^. O’Keefe and colleagues recorded normal mEPSC frequency in the hippocampus at P14 and P28, developmental periods when we observed the most significant decreases in NAc mEPSC frequency in FIRE mice relative to controls (Figure 1). O’Keefe and colleagues also observed minimal differences in transcriptome of P14 cortical neurons and astrocytes in the absence of microglia, while we found prominent difference in proteins related to synapse function, as well as significant increases in Gfap and other astrocytic proteins via NAc tissue proteomics of P22 FIRE mice and controls (Figure 2, S4, Table S1). Finally, O’Keefe and colleagues measured normal coherence of hippocampal and cortical oscillatory activity in adult FIRE mice, whereas we found significant anxiety-like phenotypes, impaired avoidance learning, and perturbed behavior-associated activity patterns in the NAc of adult FIRE mice (Figure 3,4). These divergent findings strongly support the theory that microglia play region-specific roles in synaptic development and refinement. Moreover, they may highlight critical regional differences in capacity of other CNS cells to compensate for perturbations in microglial function. Critically, microglia have been implicated in the etiology of psychiatric disorders, as they are sensitive to environmental risk factors such as chronic stress, severe early life infections, and pollution and toxin exposure^63^. There is also evidence that microglia mitigate the effects of chronic stress on neural circuit function and behavior^63,64^ and perturbed microglial function increases vulnerability to anxiety-like behaviors^65,66^. Our findings, together with those of O’Keefe et al., suggest that brain regions that are particularly relevant for psychiatric illness, such as the mesolimbic system, may be uniquely vulnerable to perturbed circuit development as a result of insults that impact microglial function.

Genetically-induced absence of microglia provides a valuable tool to easily study the effects of microglia on brain development, as microglia never form at any point throughout the lifespan. Moreover, this approach has advantages over drug-, toxin-, and CreER/tamoxifen-induced depletion, which could lead to off-target effects induced by administered substances as well as the death and clearance from the tissue of the existing microglia. However, our genetic approach lacks the temporal specificity required to reveal how microglia may be changing their functional roles at different stages of development. In addition, both genetic and drug-induced approaches lack regional selectivity and eliminate microglia across the entire brain. As tools improve to induce focal elimination of select populations of microglia, it will be important to compare the roles of local microglia within the NAc with the roles of microglia in long-range input and output structures. These studies will be crucial as we continue to identify region and subregion-specific differences in microglial phenotype and function^15,45,67^.

## Methods

### Animals

Male and female mice were used for all experiments. *Csf1r* ^ΔFIRE/WT^ mice^19^ were obtained from the Pridans lab at the University of Edinburgh and crossed with wild-type CBA mice (Jackson Laboratories #000656). Resulting *Csf1r* ^ΔFIRE/WT^ mice were crossed to produce *Csf1r* ^ΔFIRE/ΔFIRE^ (KO), *Csf1r* ^ΔFIRE/WT^ (HET), and *Csf1r* ^WT/WT^ (WT) littermates. For some experiments, HET mice were included in the microglia-positive control group, as these mice have no detectable difference in microglia number compared to WT controls^19^. Mice were housed on a 12-hour light cycle (lights on 6am-6pm). All procedures followed animal care guidelines approved by the University of California, Los Angeles Chancellor’s Animal Research Committee.

### Immunohistochemistry

Mice were anesthetized with isoflurane and transcardially perfused with phosphate-buffered saline (PBS) followed by 4% paraformaldehyde (PFA). For c-Fos experiments, mice were perfused 60 minutes after the end of behavioral analyses. Brains were post-fixed in PFA for 4-24 hours and transferred to PBS + 0.01% sodium azide (PBS/NaN_3_). 60µm brain sections were prepared on a vibratome or cryostat. Sections were washed 3 x 10 min in PBS and transferred to a blocking solution (2% NDS, 3% BSA, 0.3% Triton-X in PBS) for 2 hours. Sections were incubated overnight at 4°C in blocking solution containing primary antibody (Rabbit Anti-GFAP (Agilent #Z0334, 1:500), Mouse Anti-S100β (Sigma-Aldrich #S2532, 1:500), Mouse Anti-NeuN (EMD Millipore #MAB377, 1:500), Chicken Anti-TH (Aves #TYH, 1:500), Rabbit Anti-c-Fos (Synaptic Systems #226008, 1:1000)), or 4 nights for c-Fos experiments. After 3 x 10 min washes in PBS, sections were incubated in PBS containing secondary antibody (Alexa 488 Donkey Anti-Rabbit (Jackson ImmunoResearch #711-545-152) or Anti-Chicken (#703-545-155), Alexa 594 Donkey Anti-Chicken (#703-585-155), Alexa 647 Donkey Anti-Mouse (#715-605-151), all 1:1000), 2% NDS, and 3% BSA for 2 hours at room temperature. Sections were then washed in PBS, stained with DAPI (1:4000), and mounted using Fluoromount G. Images containing NAc were obtained at 20x using a Zeiss Axio Imager 2 fitted with Apotome 2, at 20x on a Leica Stellaris Confocal microscope, or at 5x on a Leica DM6 B scanning microscope (for c-Fos imaging). 20x S100β and GFAP images were collected as z-stacks with a 0.75μm interval, and 20x IBA1 images were collected as z-stacks with a 1.5μm interval.

To calculate microglia and astrocyte density, IBA1+ microglia or S100B+ or GFAP+ astrocytes were manually counted in two ROIs per 20x image, and each data point represents the average cell density in NAc across 2-3 brain sections per animal. Fos+ neurons were detected using our deep-learning based cell-detection algorithm DeepCOUNT^68^, with minor modifications for use in 2D histological sections.

### Acute Brain Slice Preparation

Mice were anesthetized with isoflurane. Heads were removed and placed in ice-cold NMDG cutting solution, consisting of (in mM) 92 NMDG, 20 HEPES, 30 NaHCO_3_, 1.2 NaH_2_PO_4_, 2.5 KCl, 5 Na-Ascorbate, 3 Na-Pyruvate, 2 Thiourea, 10 MgSO_4_, 0.5 CaCl_2_, 25 Glucose (adjusted to pH 7.4 with HCl). Brains were rapidly dissected and 230um sections containing NAc were obtained using a Leica VT1200S Vibratome in NMDG solution. Brain sections were transferred to NMDG solution at 34°C and incubated for approximately 5 minutes prior to transfer to room-temperature ACSF consisting of 125 NaCl, 2.5 KCl, 1.25 NaH_2_PO_4_, 1 MgCl_2_, 26 NaHCO_3_, 11 Glucose, 2.4 CaCl_2_. Sections were allowed to recover for at least 45 minutes prior to recording.

### Slice Electrophysiology

NAc neurons were visualized under infrared-differential interference contrast optics. NAc was identified by the presence of the anterior commissure. All recordings were performed at room temperature. Data were collected in pClamp 10 or 11 using an Axon Instruments Multiclamp 700B amplifier and Digidata 1440A or 1550B digitizer. Series resistance was regularly monitored, was left uncompensated, and did not change by more than 20% across a recording. For measurements of excitatory currents, whole-cell voltage clamp recordings were obtained using borosilicate glass pipettes (3-7mOhm) filled with internal solution consisting of 117 CsMS, 20 HEPES, 0.4 EGTA, 2.8 NaCl, 5 TEA-Cl, 4 Na_2_-ATP, and 0.4 Na-GTP, adjusted to pH 7.3 using CsOH (280-290mOsm). Miniature excitatory postsynaptic currents (mEPSCs) were obtained at - 70mV. Cells were patched in ACSF containing 100uM picrotoxin, then 1uM tetrodotoxin was washed in prior to mEPSC recordings. Data were analyzed off-line using Clampfit (Molecular Devices), Origin (OriginLab), Mini analysis (Synaptosoft), and PRISM (GraphPad) software. Input resistance and membrane capacitance were calculated from a 2 mV hyperpolarizing step from a holding potential of −70 mV. mEPSC frequency (>5 pA amplitude, <1 ms rise time) was quantified by monitoring activity during continuous recording for at least 8 min. For measurements of miniature inhibitory postsynaptic currents (mIPSC), a high-chloride internal solution was used consisting of 105 CsCl_2_, 10 EGTA (CsOH), 20 TEA-Cl, 20 Hepes, 1 MgCl_2_, 2 Na_2_-ATP and 0.2 Na-GTP, adjusted to a pH at 7.3 (290 mOsm). Cells were patched in ACSF, then 1uM tetrodotoxin and 5uM CNQX was washed in prior to mIPSC recordings. Data were analyzed off-line using Clampfit (Molecular Devices), Origin (OriginLab), Mini analysis (Synaptosoft), and PRISM (GraphPad) software. Input resistance and membrane capacitance were calculated from a 2 mV hyperpolarizing step from a holding potential of −70 mV. mIPSC frequency (>5 pA amplitude, <1 ms rise time) was quantified by monitoring activity during continuous recording for at least 8 min.

Recordings involving electrical stimulation were obtained by positioning a bipolar stimulating electrode in NAc 100-200um dorsal to the cell of interest. Recordings were performed in ACSF containing 100uM picrotoxin. Traces represent the average of 10 sweeps per neuron, with 10 seconds separating each sweep. Paired-pulse ratio (PPR) recordings were obtained by delivering two 1ms pulses (15-100uA) through the stimulating electrode at intervals of 20, 50, 100, and 200ms. AMPA/NMDA ratio recordings were obtained by delivering a 1ms electrical stimulation while holding the neuron at -70mV to obtain the AMPA current, then slowly increasing the holding potential to 40mV and performing the same stimulation to obtain the NMDA current. The value for the NMDA current was taken as the value 50ms after the delivery of the stimulation, as the contribution of AMPA receptors to the current at this point is minimal due to differences in channel kinetics.

PPR and AMPA/NMDA recordings were quantified in python. All traces from a cell were averaged, and the peak current value from this trace was calculated after subtracting the holding current. PPR was calculated as the ratio between the peaks of the second and first stimulations. AMPA/NMDA ratio was calculated as the ratio between the peak of the AMPA current at -70mV and the current value 50ms after the stimulation at 40mV.

### Tissue proteomics

*Csf1r* ^ΔFIRE/ΔFIRE^ and *Csf1r* ^WT/WT^ mice age P20-22 were anesthetized with isoflurane and perfused transcardially with 10 mL of oxygenated, ice-cold NMDG solution (described above for preparation of acute brain slices for electrophysiology recordings). Brains were then rapidly dissected and coronal forebrain sections (300 µm thick) were prepared using a vibratome in ice-cold NMDG solution bubbled continuously with 95% O_2_/5% CO_2_. After sectioning, slices remained in ice-cold solution, oxygenated NMDG solution and were transferred one at a time to a glass dissecting surface under a stereoscope maintained at 4°C. NAc were rapidly microdissected, minced, and transferred to eppendorf tubes containing 1 mL of 1M PBS. Samples were centrifuged at 3000 rpm, supernatant was removed, and tissues were stored at -80°C until further processing.

All samples were resuspended in equal volume of 8M Urea and 100mM Tris-Cl (pH 8). This was followed by reduction with TCEP and alkylation with IAA, followed by protein clean up with SP3 beads. Overnight digestion was done with trypsin and lysC enzymes. The peptides were cleaned using the SP3 protocol^69^, followed by elution in 2% DMSO. Samples were dried using a speed vacuum, and the dried peptides were resuspended in 5% formic acid before being sent for liquid chromatography with tandem mass spectrometry (LC-MS/MS).

### Quantitative proteomics analysis

MaxQuant software was used for peptide identification^70^. These algorithms use correlational analyses and graph theory to measure multiple mass measurements, and then integrate across these measurements and correct for linear and nonlinear mass offsets. Raw intensity data from each sample was batch corrected using a quantile normalization followed by a median centering procedure. This procedure uses an iterative algorithm to find the point that minimizes Euclidean distance to all features in the dataset and adjusts the distribution from each batch to this point. The intensity data of each identified protein log2 normalized for analysis. To be included in the analysis, a protein needed to be detected in 75% or more of the samples for each genotype. Fold changes were calculated as the ratio of the KO signal relative to the WT signal. Statistical significance was assessed using unpaired t-tests with an alpha-level of 0.05. Process enrichment analysis of the proteins up-and down-regulated was conducted in Metascape^41^. Synaptic protein enrichment analysis was conducted using SynGo^42^.

### Platform-Mediated Avoidance

All behavior experiments were performed on adult mice between 2.5 and 4 months of age. Mice were handled for at least 10 minutes across 2-3 days prior to all behavioral experiments. For platform-mediated avoidance (PMA), mice were placed in an operant chamber with a shock floor. A quarter of the floor was covered by a plexiglass platform which prevented transmission of the shock. Two scented odor pods containing peanut butter or almond, coconut, or vanilla extract were placed beneath the shock floor to promote exploration. Following 80 seconds to explore the chamber, mice received 3 baseline tones (4kHz, 75dB, 30 seconds) separated by a randomized interval of 80-150 seconds. Mice then received 9 tone-shock pairings, with each tone co-terminating with a 13mA, 2 second foot shock. The following day, mice were placed in the same chamber and received 6 tones with no shocks. Videos of behavior were obtained using PointGray Chameleon 3 cameras (Teledyne FLIR) and analyzed using DeepLabCut^71^ and BehaviorDEPOT^53^ as previously described.

### Open Field Test & Elevated Zero Maze

For open field behavior, mice were placed in an open arena (50cm x 50cm) with 30cm walls, and behavior was recorded for 10 minutes. The center was defined as a 30cm x 30cm square in the middle of the arena. For elevated zero maze, mice were placed on a custom built apparatus 24 inches in diameter, and behavior was recorded for 10 minutes. These assays were analyzed using Biobserve Viewer software.

### Shock Sensitivity

Mice were placed in an operant chamber with a shock floor. Mice received a 2 second foot shock every 20 seconds, starting at a 0.02mA intensity and increasing in increments of 0.02mA until a vocalization response was elicited, defined as the mouse emitting an audible sound. A blinded experimenter documented the first instance of scurry, dart, and vocalization for each mouse. Scurry was defined as a slow, backward movement with shuffling of the paws. Dart was defined as a more rapid forward movement at some point during the shock period.

### Stereotaxic Surgery

Mice were anesthetized in 5% isoflurane until loss of righting reflex, and the hair was cleared from the scalp using a small razor. Mice were transferred to a stereotaxic surgical apparatus and received 2% isoflurane to maintain anesthesia. Body temperature was maintained at 40°C using a water-based heating system. The scalp was cleaned with 3 alternating swabs of betadine and 70% ethanol, and a small incision was made in the scalp over the region of interest. Carprofen (50mg/kg) was delivered subcutaneously. A small hole was drilled over the injection site, and a hamilton syringe was used to deliver 400nL of AAV-syn-jGCaMP7f-WPRE (1.80*10^13^ vg/mL, diluted 1:1 in 0.9% NaCl solution, Addgene #104488) into right NAc (AP +1.3, ML +0.65, DV - 4.6). A 400um optic fiber was then implanted at AP +1.3, ML +0.65, DV -4.55 and sealed in place using metabond. Mice were allowed to wake up under warmed conditions, and carprofen (50mg/kg) was delivered subcutaneously each day for at least two days following the surgery. Some knockout mice did not survive post-surgery.

### Fiber Photometry

Following at least 3 weeks for viral expression, mice underwent PMA as described above, but with 12 tone-shock pairings on training day and 9 tone deliveries on retrieval day to increase the number of trials over which average traces could be obtained from each mouse. Mice were handled and habituated to the fiber attachment for at least 3 days prior to recording. A TDT RZ10x processor in combination with the TDT Synapse software was used to record a 405nm isosbestic channel and the 465nm signal channel while mice performed the PMA task. TTL pulses marking the start and end of each tone were used to align behavior and photometry data. Data were analyzed as previously described by fitting a curve to the 405nm isosbestic channel using the polyfit function in MATLAB and subtracting this from the 465nm signal channel. Z scores were calculated based on the mean and SD of a baseline period of 10 seconds prior to event onset for shock responses and -20 to -15 seconds for platform entry responses. Error bars in photometry traces are based on SEM of animal averages. For visualization, traces were smoothed by averaging values across each 0.5 second period.

## Acknowledgments

We thank the UCLA Proteome Research Center and J. Wohlschlegel and Y. Jami-Alahmadi for their assistance with tissue proteomics. This work was supported by R01MH127214 (L.A.D.), Brain and Behavior Research Foundation NARSAD Young Investigator Award (L.M.D.), Brain Research Foundation Seed Grant (L.M.D.), UCLA David Geffen School of Medicine Seed Grant (L.A.D., L.M.D), F30MH134633 (M.W.G.), T32GM008042 (M.W.G.), T32NS048004 (M.W.G.).

## Author Contributions

Conceptualization: M.W.G., F.E., E.N.M., L.A.D. and L.M.D; Methodology: M.W.G, F.E., E.N.M., M.S.C., S.V.B., R.R., C.P., L.A.D. and L.M.D; Investigation: M.W.G., F.E., E.N.M., M.S.C., M.S.C., S.V.B., J.P.R., A.S.E., M.H., A.Q., Formal Analysis: M.W.G., F.E., E.N.M.; Writing – Original Draft: M.W.G., F.E., L.A.D. and L.M.D; Writing – Review & Editing: M.W.G., F.E., L.A.D. and L.M.D; Supervision: L.A.D. & L.M.D; Funding Acquisition: M.W.G., F.E., L.A.D., & L.M.D.

---

**Figure S1:**
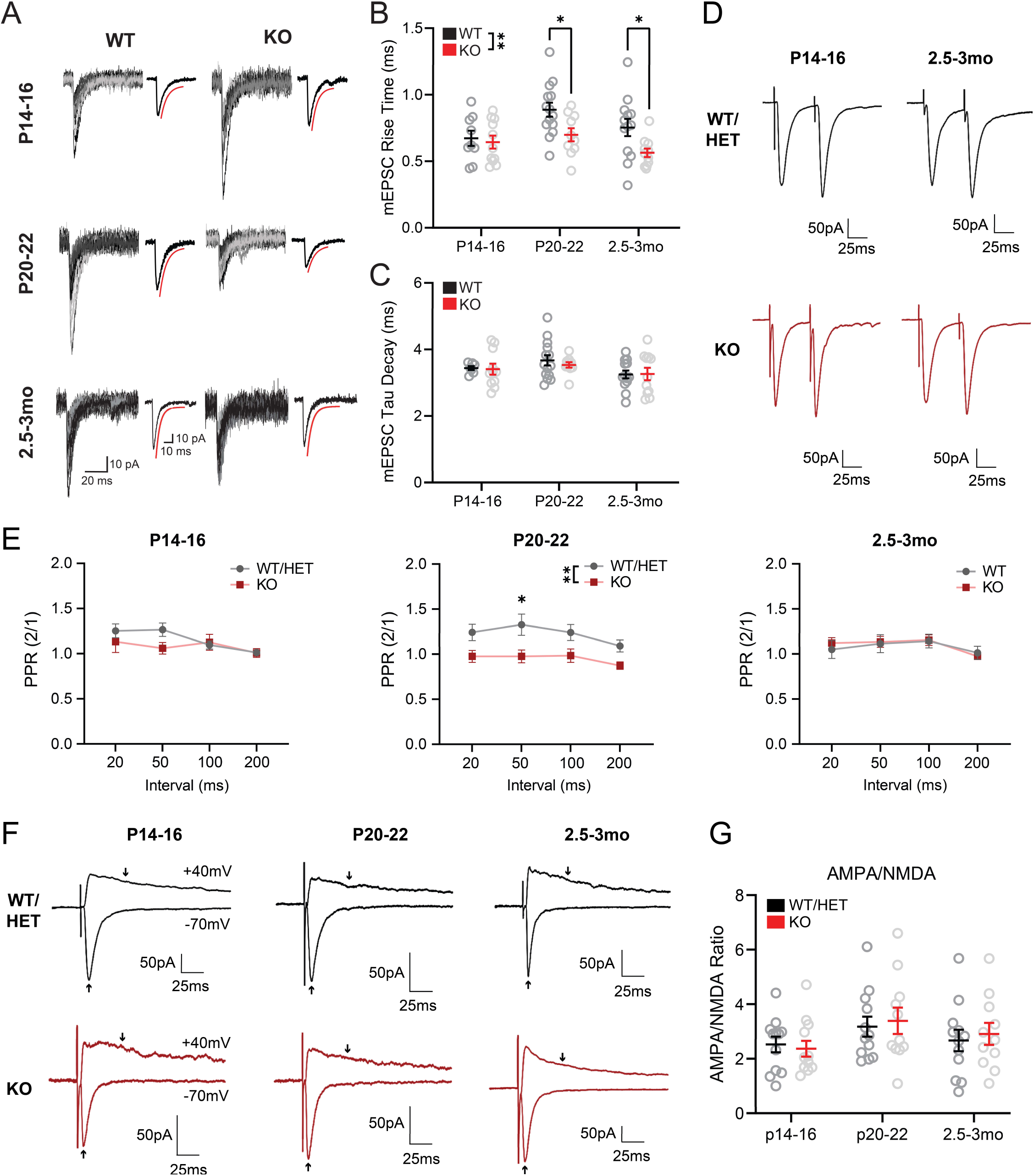
Additional electrophysiological measures of excitatory synaptic development in the presence or absence of microglia. (A) Overlay of 10 to 15 single mEPSC events in WT and KO mice at p14-16, p20-22 and 2.5-3mo. The average trace for each condition is represented on the right, and the fit exponential curve is shown in red. (B) mEPSC rise time in NAc of WT and KO mice at ages with a sufficient number of events for analysis of kinetics. F_age_(2,62)=4.268, *P* =0.0183; F_genotype_(1,62)=9.472, *P* =0.0031; F_age x genotype_(2,62)=1.390, *P* =0.2568. (C) mEPSC Tau decay constant in NAc of WT and KO mice. F_age_(2,60)=3.009, *P* =0.0569; F_genotype_(1,60)=0.1783, *P* =0.6743; F_age x genotype_(2,60)=0.1469, *P* =0.8637. Two-way ANOVA with Šídák’s multiple comparisons test. P14-16: WT *n*=9(6), KO *n*=11(5); P20-22: WT *n*=14(7), KO *n*=10(6); 2.5-3mo: WT *n*=13(7), KO *n*=11(6). (D) Representative traces of evoked paired-pulse ratio (PPR) recordings with a 50ms interstimulus interval in NAc knockout and control mice at P14-16 and 2.5-3mo. Cells were recorded while voltage clamping at -70mV in the presence of 100µM picrotoxin. (E) Paired-pulse ratio from interstimulus intervals ranging from 20 to 200ms. P14-16: F_Interval_(3,69)=5.018, *P* =0.0033; F_Genotype_(1,23)=0.8535, *P* =0.3652; F_Interval x Genotype_(3,69)=2.294, *P* =0.0855. P20-22: F_Interval_(3,60)=5.807, *P* =0.0240; F_Genotype_(1,20)=5.962, *P* =0.0015; F_Interval x Genotype_(3,60)=0.8502, *P* =0.4720. 2.5-3mo: F_Interval_(3,69)=8.863, *P* <0.0001; F_Genotype_(1,23)=0.02861, *P* =0.8672; F_Interval x Genotype_(3,69)=1.001, *P* =0.3976. Two-way RM ANOVA with Šídák’s multiple comparisons test. P14-16: WT/HET *n*=13(4), KO *n*=12(3); P20-22: WT/HET *n*=13(4), KO *n*=12(4); 2.5-3mo: WT *n*=12(4), KO *n*=13(4). (F) Representative traces of evoked AMPA/NMDA recordings in knockout and control mice. Cells were recorded while voltage clamping at -70mV or +40mV in the presence of 100µM picrotoxin. Arrows indicate timepoints used for current estimation: the NMDA current was taken as the current value from the +40mV stimulation 50 ms after stimulus onset, and the AMPA current was taken as the peak amplitude of the evoked current when held at -70mV. (G) AMPA/NMDA ratio in NAc of KO and control mice at P14-16, P20-22, and 2.5-3mo. F_Age_(2,64)=2.562, *P* =0.0850; F_Genotype_(1,64)=0.1112, *P* =0.7399; F_Age x Genotype_(2,64)=0.1802, *P* =0.8355. Two-way ANOVA. P14-16: WT/HET *n*=12(4), KO *n*=12(3); P20-22: WT/HET *n*=12(5), KO *n*=11(4); 2.5-3mo: WT *n*=12(4), KO *n*=11(4). Error bars represent mean ± SEM. \**P* <0.05. \*\**P* <0.01.

**Figure S2:**
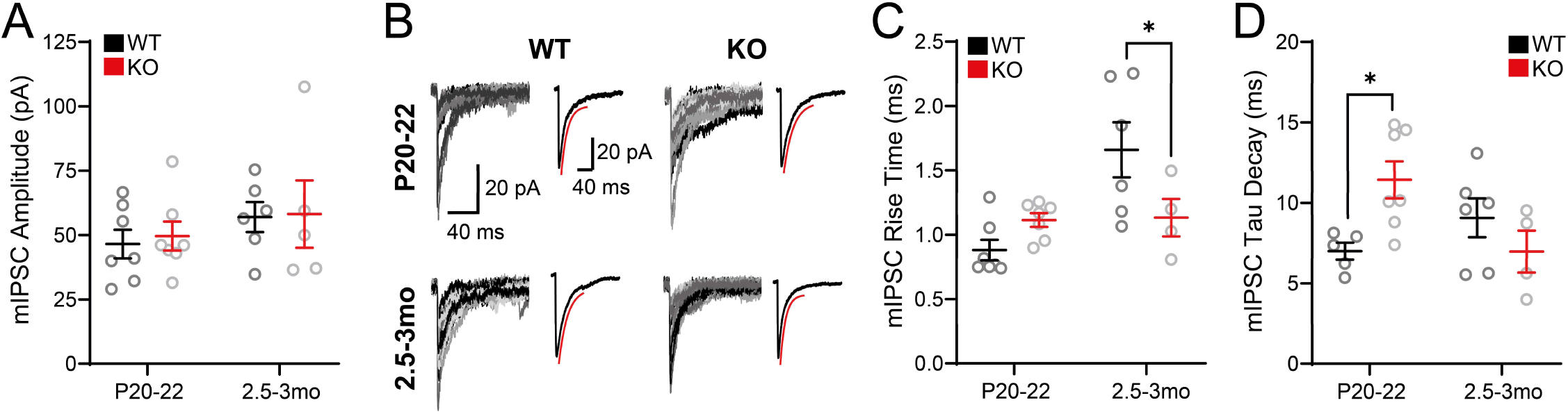
Additional electrophysiological measures of inhibitory synaptic development in the presence or absence of microglia. (A) mIPSC amplitude in NAc of WT and KO mice at P20-22 and 2.5-3mo. F_age_(1,21)=1.636, *P* =0.2148; F_genotype_(1,21)=0.08115, *P* =0.7785; F_age x genotype_(1,21)=0.0169, *P* =0.8978. (B) Overlay of 10 to 15 single mIPSC events in both WT and KO mice at P20-22 and 2.5-3mo. The average trace for each condition is represented on the right, and the fit exponential curve is shown in red. (C-D) mIPSC rise time (C) and Tau decay constant (D) in NAc of WT and KO mice at P20-22 and 2.5-3mo. Rise time: F_age_(1,20)=8.977, *P* =0.0071; F_genotype_(1,20)=1.225, *P* =0.2815; F_age x genotype_(1,20)=8.159, *P* =0.0098; Tau Decay: F_age_(1,18)=1.074, *P* =0.3137; F_genotype_(1,18)=1.031, *P* =0.3233; F_age x genotype_(1,18)=8.058, *P* =0.0109. Two-way ANOVA with Šídák’s multiple comparisons test. P20-22: WT n=7(3), KO n=7(3); 2.5-3mo: WT n=6(3), KO n=5(3). \**P* <0.05. Error bars represent mean ± SEM.

**Figure S3:**
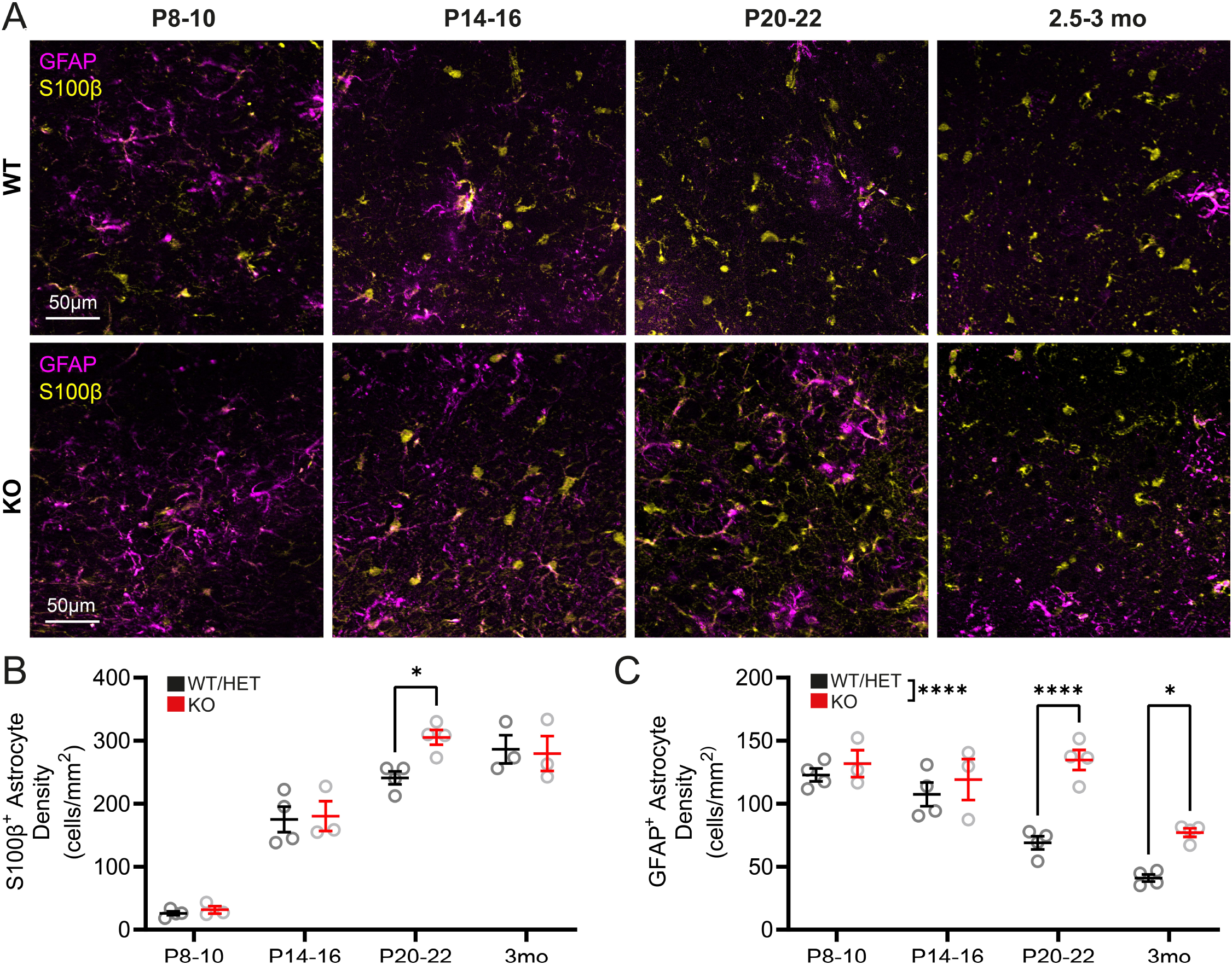
Absence of microglia alters normal developmental trajectories of astrocyte markers in NAc. (A) Representative images of S100β_+_ and GFAP_+_ astrocytes in NAc of WT and KO mice at p8-10, p14-16, p20-22 and 2.5-3mo. Zoom panels represent the area with a dashed outline. (B) S100β_+_ astrocyte density in NAc of WT and KO mice across postnatal development. F_age_(3,20)=99.74, *P* <0.0001; F_genotype_(1,20)=2.092, *P* =0.1635; F_age x genotype_(3,20)=2.004, *P* =0.1458, Two-way ANOVA with Šídák’s multiple comparisons test. P8-10: WT *n*=3, KO *n*=3; P14-16: WT *n*=4, KO *n*=3; P20-22: WT *n*=3, KO *n*=4; 2.5-3mo: WT *n*=3, KO *n*=3. (C) GFAP_+_ astrocyte density in KO and control mice across postnatal development. F_age_(3,22)=29.29, *P* <0.0001; F_genotype_(1,22)=30.90, *P* <0.0001; F_age x genotype_(3,22)=5.798, *P* =0.0045; Two-way ANOVA with Šídák’s multiple comparisons test. P8-10: WT *n*=4, KO *n*=3; P14-16: WT *n*=4, KO *n*=3; P20-22: WT *n*=3, KO *n*=4; 2.5-3mo: WT/HET *n*=4, KO *n*=4. \**P* <0.05, \*\*\*\**P* <0.0001. Error bars represent mean ± SEM.

**Figure S4.**
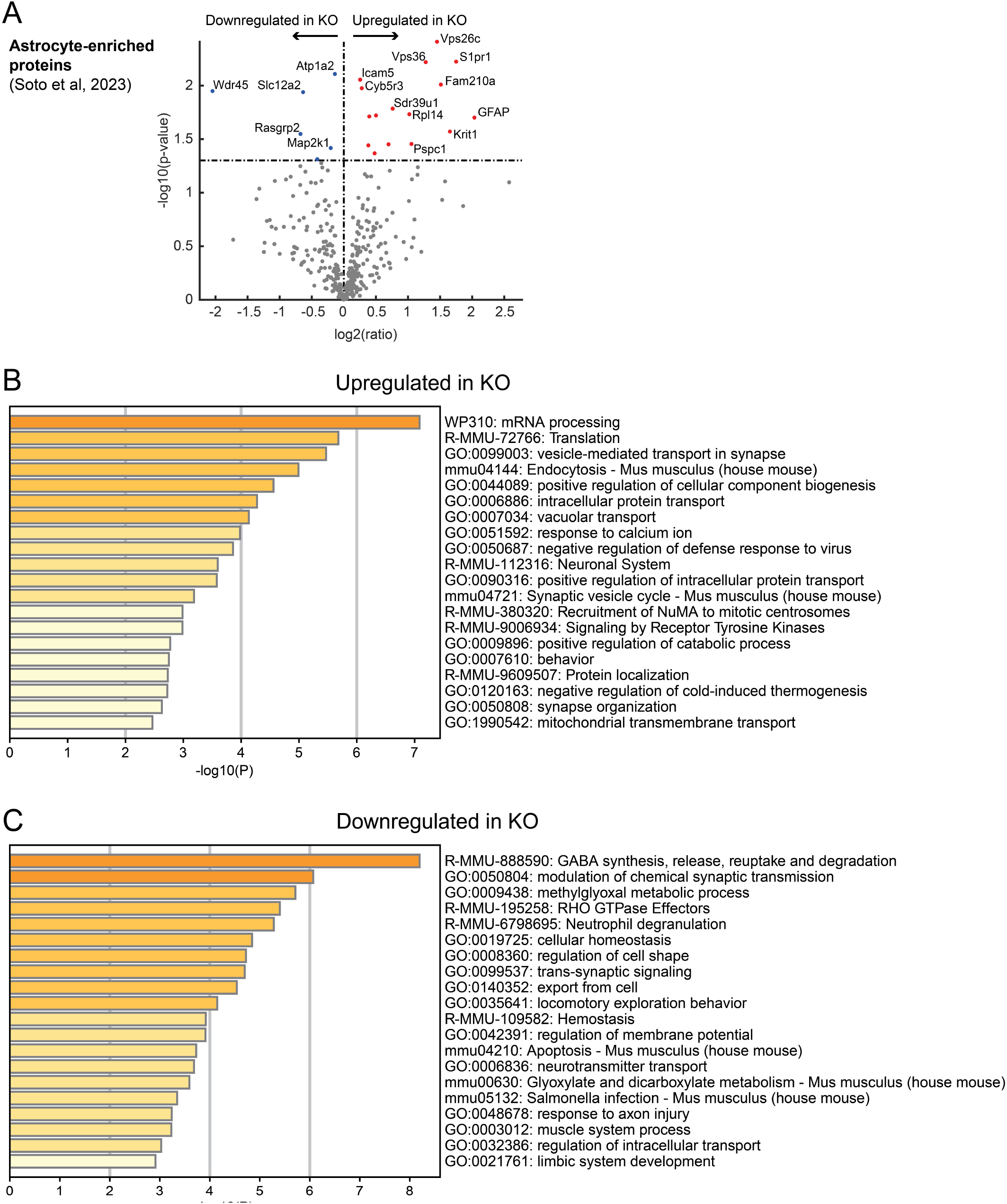
Additional analysis of proteomic dataset. (A) Volcano plot of protein expression in WT vs KO mice in 342 proteins shown to be enriched/unique for astrocytes compared to neurons in Soto et. al. 2023. (B-C) Complete list of top 20 pathways upregulated (B) and downregulated (C) in KO mice upon Metascape analysis of differentially expressed proteins from proteomic data.

**Figure S5.**
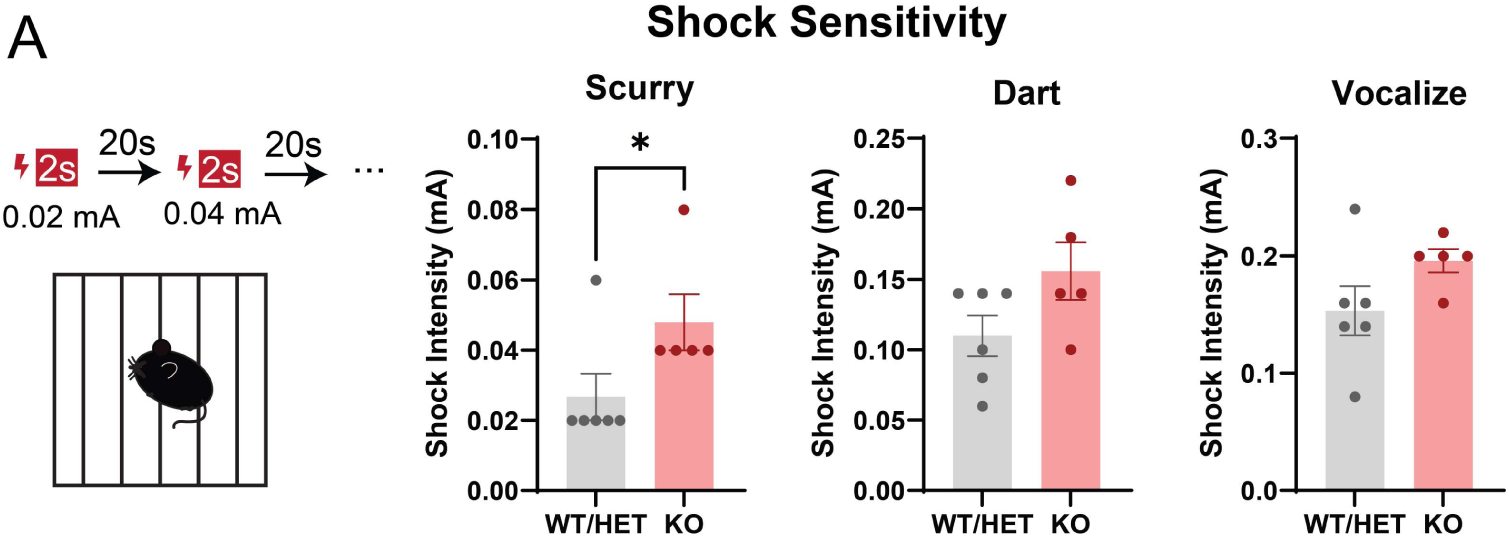
(A) Minimum shock intensity required to elicit scurry, dart, and vocalization responses in knockout and control mice. Mann-Whitney test, WT/HET: *n*=6, KO: *n*=5. \**P* <0.05. Error bars represent mean ± SEM.

## References

1. Squarzoni, P. et al. Microglia Modulate Wiring of the Embryonic Forebrain. Cell Rep. 8, 1271–1279 (2014).

2. Hammond, T. R., Robinton, D. & Stevens, B. Microglia and the Brain: Complementary Partners in Development and Disease. Annu. Rev. Cell Dev. Biol. 34, 523–544 (2018).

3. Frost, J. L. & Schafer, D. P. Microglia: Architects of the Developing Nervous System. Trends Cell Biol. 26, 587–597 (2016).

4. Lukens, J. R. & Eyo, U. B. Microglia and neurodevelopmental disorders. Annu. Rev. Neurosci. 45, 425–445 (2022).

5. Reemst, K., Noctor, S. C., Lucassen, P. J. & Hol, E. M. The Indispensable Roles of Microglia and Astrocytes during Brain Development. Front. Hum. Neurosci. 10, (2016).

6. Blagburn-Blanco, S. V., Chappell, M. S., De Biase, L. M. & DeNardo, L. A. Synapse-specific roles for microglia in development: New horizons in the prefrontal cortex. Front. Mol. Neurosci. 15, (2022).

7. Gallo, N. B., Berisha, A. & Van Aelst, L. Microglia regulate chandelier cell axo-axonic synaptogenesis. Proc. Natl. Acad. Sci. 119, e2114476119 (2022).

8. Miyamoto, A. et al. Microglia contact induces synapse formation in developing somatosensory cortex. Nat. Commun. 7, 12540 (2016).

9. Parkhurst, C. N. et al. Microglia Promote Learning-Dependent Synapse Formation through Brain-Derived Neurotrophic Factor. Cell 155, 1596–1609 (2013).

10. Floresco, S. B. The nucleus accumbens: an interface between cognition, emotion, and action. Annu. Rev. Psychol. 66, 25–52 (2015).

11. Hanson, J. L., Williams, A. V., Bangasser, D. A. & Peña, C. J. Impact of Early Life Stress on Reward Circuit Function and Regulation. Front. Psychiatry 12, 744690 (2021).

12. Birnie, M. T. et al. Plasticity of the Reward Circuitry After Early-Life Adversity: Mechanisms and Significance. Biol. Psychiatry 87, 875–884 (2020).

13. Luo, L. Architectures of neuronal circuits. Science 373, eabg7285 (2021).

14. de Wit, J., Hong, W., Luo, L. & Ghosh, A. Role of leucine-rich repeat proteins in the development and function of neural circuits. Annu. Rev. Cell Dev. Biol. 27, 697–729 (2011).

15. Hope, K. T., Hawes, I. A., Moca, E. N., Bonci, A. & De Biase, L. M. Maturation of the microglial population varies across mesolimbic nuclei. Eur. J. Neurosci. 52, 3689–3709 (2020).

16. Kopec, A. M., Smith, C. J., Ayre, N. R., Sweat, S. C. & Bilbo, S. D. Microglial dopamine receptor elimination defines sex-specific nucleus accumbens development and social behavior in adolescent rats. Nat. Commun. 9, 3769 (2018).

17. Peixoto, R. T., Wang, W., Croney, D. M., Kozorovitskiy, Y. & Sabatini, B. L. Early hyperactivity and precocious maturation of corticostriatal circuits in Shank3B−/− mice. Nat. Neurosci. 19, 716–724 (2016).

18. Krajeski, R. N., Macey-Dare, A., van Heusden, F., Ebrahimjee, F. & Ellender, T. J. Dynamic postnatal development of the cellular and circuit properties of striatal D1 and D2 spiny projection neurons. J. Physiol. 597, 5265–5293 (2019).

19. Rojo, R. et al. Deletion of a Csf1r enhancer selectively impacts CSF1R expression and development of tissue macrophage populations. Nat. Commun. 10, 3215 (2019).

20. Hoshiko, M., Arnoux, I., Avignone, E., Yamamoto, N. & Audinat, E. Deficiency of the Microglial Receptor CX3CR1 Impairs Postnatal Functional Development of Thalamocortical Synapses in the Barrel Cortex. J. Neurosci. 32, 15106–15111 (2012).

21. Favuzzi, E. et al. GABA-receptive microglia selectively sculpt developing inhibitory circuits. Cell 184, 4048–4063.e32 (2021).

22. Vainchtein, I. D. et al. Astrocyte-derived interleukin-33 promotes microglial synapse engulfment and neural circuit development. Science 359, 1269–1273 (2018).

23. Liddelow, S. A. et al. Neurotoxic reactive astrocytes are induced by activated microglia. Nature 541, 481–487 (2017).

24. Bohlen, C. J. et al. Diverse Requirements for Microglial Survival, Specification, and Function Revealed by Defined-Medium Cultures. Neuron 94, 759–773.e8 (2017).

25. VanRyzin, J. W. et al. Microglial Phagocytosis of Newborn Cells Is Induced by Endocannabinoids and Sculpts Sex Differences in Juvenile Rat Social Play. Neuron 102, 435–449.e6 (2019).

26. Chung, W.-S., Baldwin, K. T. & Allen, N. J. Astrocyte Regulation of Synapse Formation, Maturation, and Elimination. Cold Spring Harb. Perspect. Biol. 16, a041352 (2024).

27. Nagai, J. et al. Behaviorally consequential astrocytic regulation of neural circuits. Neuron 109, 576 (2020).

28. Tan, C. X., Burrus Lane, C. J. & Eroglu, C. Role of astrocytes in synapse formation and maturation. Curr. Top. Dev. Biol. 142, 371–407 (2021).

29. Raponi, E. et al. S100B expression defines a state in which GFAP-expressing cells lose their neural stem cell potential and acquire a more mature developmental stage. Glia 55, 165–177 (2007).

30. Du, J. et al. S100B is selectively expressed by gray matter protoplasmic astrocytes and myelinating oligodendrocytes in the developing CNS. Mol. Brain 14, 154 (2021).

31. Landry, C. F., Ivy, G. O. & Brown, I. R. Developmental expression of glial fibrillary acidic protein mRNA in the rat brain analyzed by in situ hybridization. J. Neurosci. Res. 25, 194– 203 (1990).

32. Saunders, A. et al. Molecular Diversity and Specializations among the Cells of the Adult Mouse Brain. Cell 174, 1015–1030.e16 (2018).

33. Rangaraju, S. et al. Quantitative proteomics of acutely-isolated mouse microglia identifies novel immune Alzheimer’s disease-related proteins. Mol. Neurodegener. 13, 34 (2018).

34. Südhof, T. C. Towards an Understanding of Synapse Formation. Neuron 100, 276–293 (2018).

35. Zipursky, S. L. & Sanes, J. R. Chemoaffinity revisited: dscams, protocadherins, and neural circuit assembly. Cell 143, 343–353 (2010).

36. Dewa, K.-I. & Arimura, N. Neuronal and astrocytic protein connections and associated adhesion molecules. Neurosci. Res. 187, 14–20 (2023).

37. Sugatha, J. et al. Insights into cargo sorting by SNX32 and its role in neurite outgrowth. eLife 12, e84396 (2023).

38. Telek, E., Kengyel, A. & Bugyi, B. Myosin XVI in the Nervous System. Cells 9, 1903 (2020).

39. Erreger, K., Chen, P. E., Wyllie, D. J. A. & Traynelis, S. F. Glutamate receptor gating. Crit. Rev. Neurobiol. 16, 187–224 (2004).

40. Goetz, T., Arslan, A., Wisden, W. & Wulff, P. GABA(A) receptors: structure and function in the basal ganglia. Prog. Brain Res. 160, 21–41 (2007).

41. Zhou, Y. et al. Metascape provides a biologist-oriented resource for the analysis of systems-level datasets. Nat. Commun. 10, 1523 (2019).

42. Koopmans, F. et al. SynGO: An Evidence-Based, Expert-Curated Knowledge Base for the Synapse. Neuron 103, 217–234.e4 (2019).

43. Bennett, M. L. et al. New tools for studying microglia in the mouse and human CNS. Proc. Natl. Acad. Sci. U. S. A. 113, E1738–1746 (2016).

44. Zhang, Y. et al. An RNA-sequencing transcriptome and splicing database of glia, neurons, and vascular cells of the cerebral cortex. J. Neurosci. Off. J. Soc. Neurosci. 34, 11929– 11947 (2014).

45. De Biase, L. M. et al. Local Cues Establish and Maintain Region-Specific Phenotypes of Basal Ganglia Microglia. Neuron 95, 341–356.e6 (2017).

46. Vidak, E., Javoršek, U., Vizovišek, M. & Turk, B. Cysteine Cathepsins and Their Extracellular Roles: Shaping the Microenvironment. Cells 8, 264 (2019).

47. Tran, A. P. & Silver, J. Cathepsins in neuronal plasticity. Neural Regen. Res. 16, 26–35 (2021).

48. Niemeyer, C., Matosin, N., Kaul, D., Philipsen, A. & Gassen, N. C. The Role of Cathepsins in Memory Functions and the Pathophysiology of Psychiatric Disorders. Front. Psychiatry 11, 718 (2020).

49. Fehon, R. G., McClatchey, A. I. & Bretscher, A. Organizing the cell cortex: the role of ERM proteins. Nat. Rev. Mol. Cell Biol. 11, 276–287 (2010).

50. Bravo-Rivera, C., Roman-Ortiz, C., Brignoni-Perez, E., Sotres-Bayon, F. & Quirk, G. J. Neural Structures Mediating Expression and Extinction of Platform-Mediated Avoidance. J. Neurosci. 34, 9736–9742 (2014).

51. Bravo-Rivera, C., Roman-Ortiz, C., Montesinos-Cartagena, M. & Quirk, G. J. Persistent active avoidance correlates with activity in prelimbic cortex and ventral striatum. Front. Behav. Neurosci. 9, (2015).

52. He, Z.-X., et al. A Nucleus Accumbens Tac1 Neural Circuit Regulates Avoidance Responses to Aversive Stimuli. Int. J. Mol. Sci. 24, 4346 (2023).

53. Gabriel, C. J. et al. BehaviorDEPOT is a simple, flexible tool for automated behavioral detection based on markerless pose tracking. eLife 11, e74314 (2022).

54. Klune, C. B. et al. Developmentally distinct architectures in top-down circuits. 2023.08.27.555010 Preprint at 10.1101/2023.08.27.555010 (2023).

55. Wilton, D. K., Dissing-Olesen, L. & Stevens, B. Neuron-Glia Signaling in Synapse Elimination. Annu. Rev. Neurosci. 42, 107–127 (2019).

56. de Deus, J. L., Faborode, O. S. & Nandi, S. Synaptic Pruning by Microglia: Lessons from Genetic Studies in Mice. Dev. Neurosci. 1–21 (2024) doi:10.1159/000541379.

57. Reshef, R. et al. The role of microglia and their CX3CR1 signaling in adult neurogenesis in the olfactory bulb. eLife 6, e30809 (2017).

58. Gray, D. T. et al. Extracellular matrix remodeling during aging aligns with synapse, microglia, and cognitive status. 2024.01.04.574215 Preprint at 10.1101/2024.01.04.574215 (2024).

59. Chinthalapudi, K., Rangarajan, E. S. & Izard, T. The interaction of talin with the cell membrane is essential for integrin activation and focal adhesion formation. Proc. Natl. Acad. Sci. U. S. A. 115, 10339–10344 (2018).

60. Park, Y. K. & Goda, Y. Integrins in synapse regulation. Nat. Rev. Neurosci. 17, 745–756 (2016).

61. Kato, G. et al. Microglial Contact Prevents Excess Depolarization and Rescues Neurons from Excitotoxicity. eNeuro 3, ENEURO.0004-16.2016 (2016).

62. O’Keeffe, M. et al. Typical development of synaptic and neuronal properties can proceed without microglia in the cortex and thalamus. Nat. Neurosci. 1–12 (2025) doi:10.1038/s41593-024-01833-x.

63. Hanamsagar, R. & Bilbo, S. D. Environment Matters: Microglia Function and Dysfunction in a Changing World. Curr. Opin. Neurobiol. 47, 146–155 (2017).

64. Chen, D. et al. Microglia govern the extinction of acute stress-induced anxiety-like behaviors in male mice. Nat. Commun. 15, 449 (2024).

65. Stein, D. J., Vasconcelos, M. F., Albrechet-Souza, L., Ceresér, K. M. M. & de Almeida, R. M. M. Microglial Over-Activation by Social Defeat Stress Contributes to Anxiety- and Depressive-Like Behaviors. Front. Behav. Neurosci. 11, (2017).

66. León-Rodríguez, A., Fernández-Arjona, M. del M., Grondona, J. M., Pedraza, C. & López-Ávalos, M. D. Anxiety-like behavior and microglial activation in the amygdala after acute neuroinflammation induced by microbial neuraminidase. Sci. Rep. 12, 11581 (2022).

67. Stogsdill, J. A. et al. Pyramidal neuron subtype diversity governs microglia states in the neocortex. Nature 608, 750–756 (2022).

68. Gongwer, M. W. et al. Brain-wide projections and differential encoding of prefrontal neuronal classes underlying learned and innate threat avoidance. J. Neurosci. (2023) doi:10.1523/JNEUROSCI.0697-23.2023.

69. Hughes, C. S. et al. Single-pot, solid-phase-enhanced sample preparation for proteomics experiments. Nat. Protoc. 14, 68–85 (2019).

70. Cox, J. & Mann, M. MaxQuant enables high peptide identification rates, individualized p.p.b.-range mass accuracies and proteome-wide protein quantification. Nat. Biotechnol. 26, 1367–1372 (2008).

71. Mathis, A. et al. DeepLabCut: markerless pose estimation of user-defined body parts with deep learning. Nat. Neurosci. 21, 1281–1289 (2018).

